# Novel wheat germ agglutinin-based mass cytometry cell barcoding reagent for heterogeneous, live or fixed sample

**DOI:** 10.1101/2024.11.05.618916

**Authors:** Riley T Hannan, Dayton Barker, Brendan P Cox, Colleen A Roosa, Taylor A. Harper, Michael D. Solga, Donald R. Griffin, Jeffrey M. Sturek

## Abstract

Sample multiplexing in flow cytometry is a powerful technique which allows for reduction of error, inclusion of control samples for batch effect correction, and reduction in both time and consumable usage. Current industry standard for barcoding in mass cytometry is an intracellular reagent, which requires fixation and permeabilization of sample prior to barcoding. We developed a barcode using the ubiquitous and well-tolerated membrane labeling lectin, wheat germ agglutinin. This barcode effectively labels all tested cell types, both live and fixed. We determine that barcode yields, or the ratio of debarcoded cells to total input cells, is stable in live pooled sample for at least an hour. This barcode does not show differential performance across major PBMC lineages. Thus, this universal wheat germ agglutinin-based barcode represents an advance in gentle, non-reactive cell surface barcoding for live cells.

## Introduction

Flow cytometry is a powerful method to generate per-cell data with high throughput and increasingly higher dimensionality. With higher dimensionality, experimental protocols become more complex and the risk of introducing variability between samples, referred to henceforth as sample error, builds. Sample multiplexing in flow cytometry is a technique which serves to retire some of those risks. Available dimensionality (discretized into individual channels) can be provisioned for a set of unique channel combinations, or barcodes, from which one is applied per sample. These barcoded samples are then pooled into a single tube, processed as in a typical cytometry workflow, and acquired as a single tube. The acquired data are then deconvolved back into separate samples in software post-acquisition. Sample error from operator and instrument is eliminated in all steps subsequent to pooling^1^. Also significant is the reduction in labor and consumables required during both sample preparation and acquisition. Lastly, sample multiplexing enables easy addition of known standards (anchor samples) for inter-experimental normalization^2,3^.

Doublet-discriminating barcodes trade the maximum potential amount of a barcode, or ‘plex’, for heightened stringency in the integrity of debarcoded samples. Instead of generating barcodes with every possible binary permutation of *n* channels (2^n^ barcodes), each doublet-discriminating barcode uses exactly half the available channels (n/2 channels). This arrangement allows for *n*!/(*n*/2!)^2^ barcodes, or 20 when *n*=6, and 70 when *n*=8. This system is called doublet-discriminating as any multiplet comprised of multiple barcodes will, by definition, have more than n/2 channels labeled, allowing removal during debarcoding^4^. Events which cannot be positively assigned a barcode are assigned to a pool of “Non-debarcoded” events. Additionally, each successfully debarcoded event is assigned a value in two new channels appended to debarcoded files. These two channels are generated during debarcoding and aid in further quality control of debarcoded events. We refer to those events which do not pass those quality control thresholds as “Bad” DeBarcoded events, and those which pass as “Good” DeBarcoded events. It is these “Good” DeBarcoded events which are tacitly used in every experiment utilizing doublet-discriminating barcodes, even if this process is not mentioned. This is explored more fully in Fread et al.^5^ and Zunder et al.^4^. A schematic of the whole barcoding and debarcoding workflow can be seen in Figure 1A.

**Figure 1:**
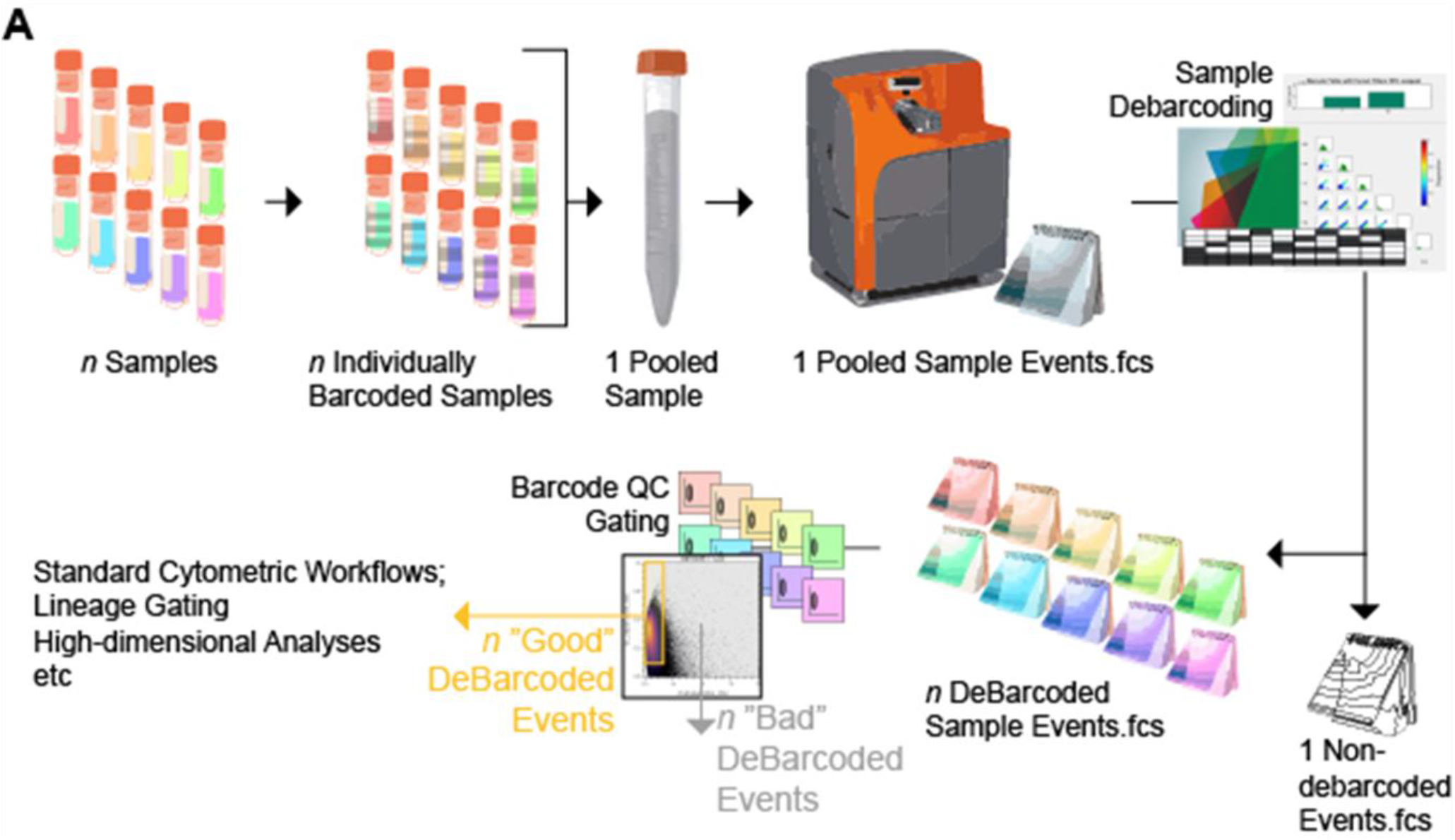
Schematic of workflow for sample barcoding and debarcoding in mass cytometry.

Standard BioTools (formerly Fluidigm) offers a doublet-discriminating barcode for mass cytometry (CyTOF), containing twenty barcodes in a *six-pick-three* arrangement. Being intracellular, the use of this barcode necessitates the fixation and permeabilization of cells. If the desired antibody panel contains fixation-sensitive antigen, staining operations must occur prior to sample barcoding and pooling. Some efforts in barcoding have therefore sought to target living cells, enabling barcoding as the first step in a workflow. The most prevalent application of live barcode is on human peripheral blood mononuclear cells (PBMCs), using anti-CD45 antibody^6–11^. Necessarily, these CD45-based barcodes are limited to the hematopoietic lineage of humans. Near-ubiquitous cell surface proteins such as MHC class I self-antigen, transmembrane transport complexes, and adhesion molecules have been used in other, more ‘universal’ live-cell barcoding reagents^8,12,13^. These reagents enable the pooled analysis of heterogeneous sample such as whole tissue, tumor, or complex cell culture assays, with the caveat of usually being species specific.

Chemical crosslinking reagents, typically thiol- or amine-reactive, are broadly applicable to all cells and have seen use in barcoding applications^4,14–17^. These reagents covalently bind to proteins on a cell surface, and have been used to barcode live cells in suspension or organoid culture^16,17^. Tellurium-maleimide (TeMal) reactive reagent has recently been made commercially available by Standard BioTools^18^. Another barcoding reagent which does not fit neatly into the affinity or reactive categories employs polystyrene nanoparticles which are internalized by cells over the course of hours^19^.

Here, we report the use of Wheat Germ Agglutinin (WGA) as a universal cell barcoding reagent for sample multiplexing in CyTOF. The isolation and identification of WGA as a glycoprotein was first reported in 1967^20^. It is a dimeric lectin with high affinity for sialic acid and N-acetylglucosamine (GlcNAc), ubiquitous residues in animal cells^21^. It rapidly gained popularity in the 1980s as a label in neuronal tracing studies^22^,and is now a common membrane-labeling tool available as a conjugate or free molecule. WGA has been used in flow cytometric analyses, including in CyTOF, as a proxy for cell size and for characterization of bacterial surface sugars ^23–25^. Its broad use as a pan-membrane marker makes it a compelling reagent for barcoding. We generated thiolated WGA (tWGA) and validated tWGA’s membrane-labeling capacity post-thiolation. Using available maleimide conjugation kits, we generated a range of monoisotopically conjugated tWGA. These metal-conjugated tWGAs were combined to generate doublet-discriminating, sample multiplexing barcodes. This barcode performs similarly to CD45 live surface barcode and palladium intracellular barcode in sample multiplexing experiments, while effectively barcoding all cell types tested. Finally, we examined the stability of tWGA barcoding in pooled sample and found tWGA debarcoding yields remain stable for at last an hour of sample pooling prior to surface staining and fixation.

## Results

### Thiolation of wheat germ agglutinin using Traut’s reagent

WGA was thiolated using 2-iminothialane (Traut’s reagent) in PBS+2mM EDTA, pH 8.0, at room temperature for one hour to generate stably thiolated WGA (tWGA), reaction schematized in Figure 2A. A range of molar excesses of Traut’s reagent were tested, with thiolation confirmed by Ellman’s assay (Supplemental Figure 1), and tWGA subsequently conjugated with AlexaFluor 647 C2 maleimide (+AF647-mal) for interrogation via microscopy (representative micrographs, Figure 2B) and quantitation of per-cell integrated fluorescence intensity (Figure 2C). Data points refer to well replicates. A tenfold molar excess of Traut’s reagent to WGA was chosen for all subsequent reactions. Staining PMBCs with the same tWGA-647 at four, twenty-five, and thirty-seven degrees Celsius revealed no differences in staining intensity by Imaging flow cytometry (Supplemental Figure 2).

**Figure 2:**
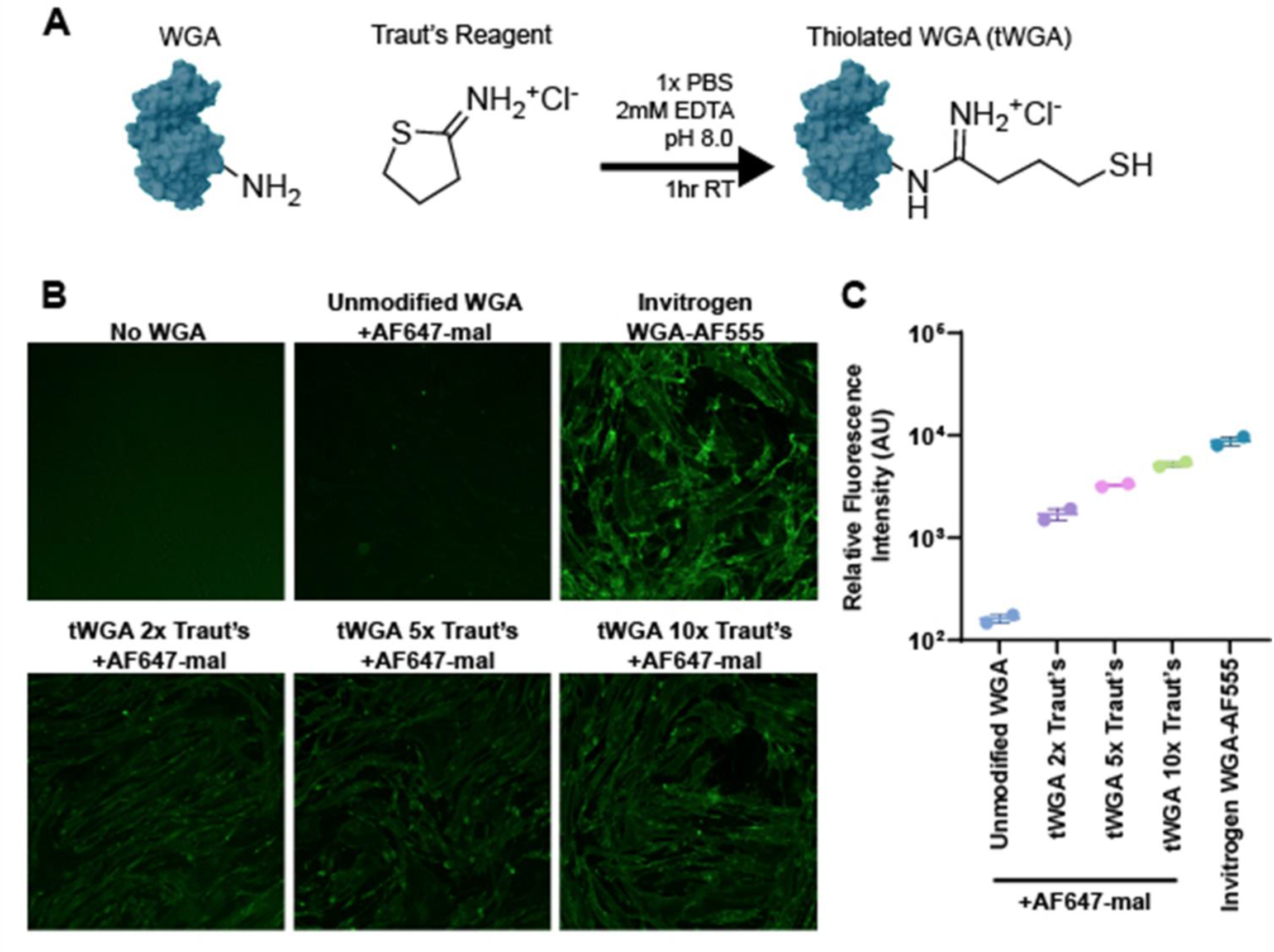
2-iminothialane (Traut’s reagent) thiolation of WGA (tWGA) enables maleimide conjugation and preserves WGA’s membrane-staining. (A) Reaction of WGA amines with Traut’s reagent in 2mM EDTA in PBS at pH 8.0, room temperature, for one hour results in a product of thiolated WGA (tWGA). (B) Representative immunofluorescence micrographs of human dermal fibroblasts labeled with different preparations of tWGA conjugated to AlexaFluor 647 C2 maleimide. (C) Quantitation of mean relative cellular fluorescence intensity per field.

### tWGA-metal conjugates stain cells from various tissues and fixations

tWGA conjugated to 89Y (tWGA-89Y) was used to stain live human PBMCs, paraformaldehyde(PFA)-fixed human PBMCs and whole blood processed with PROT1 Proteomic Stabilizer (SmartTube Inc) (Figure 3A). A range of titers was used, corresponding to standard test volumes of a 0.1mg/mL antibody. E.g. 20µg/mL equivalent to 20uL of 0.1mg/mL antibody in a 100uL test volume. Positive staining of tWGA-89Y was seen with titers of 1.25 and above. A minimal surface marker panel was included to visualize the varying intensities of tWGA-89Y across leukocyte lineages. Lymphocytes reported lower mean intensity of tWGA than myeloid lineages. PFA-fixed mouse lung digest stained with a range of tWGA-116Cd titers demonstrates positive staining at the lowest titer tested, 0.3125µg/mL. (Figure 3B). Cells stained with tWGA-89Y were added in a 1:1 ratio into the 1.25µg/mL tWGA-116Cd sample for visualization of a negative population.

**Figure 3:**
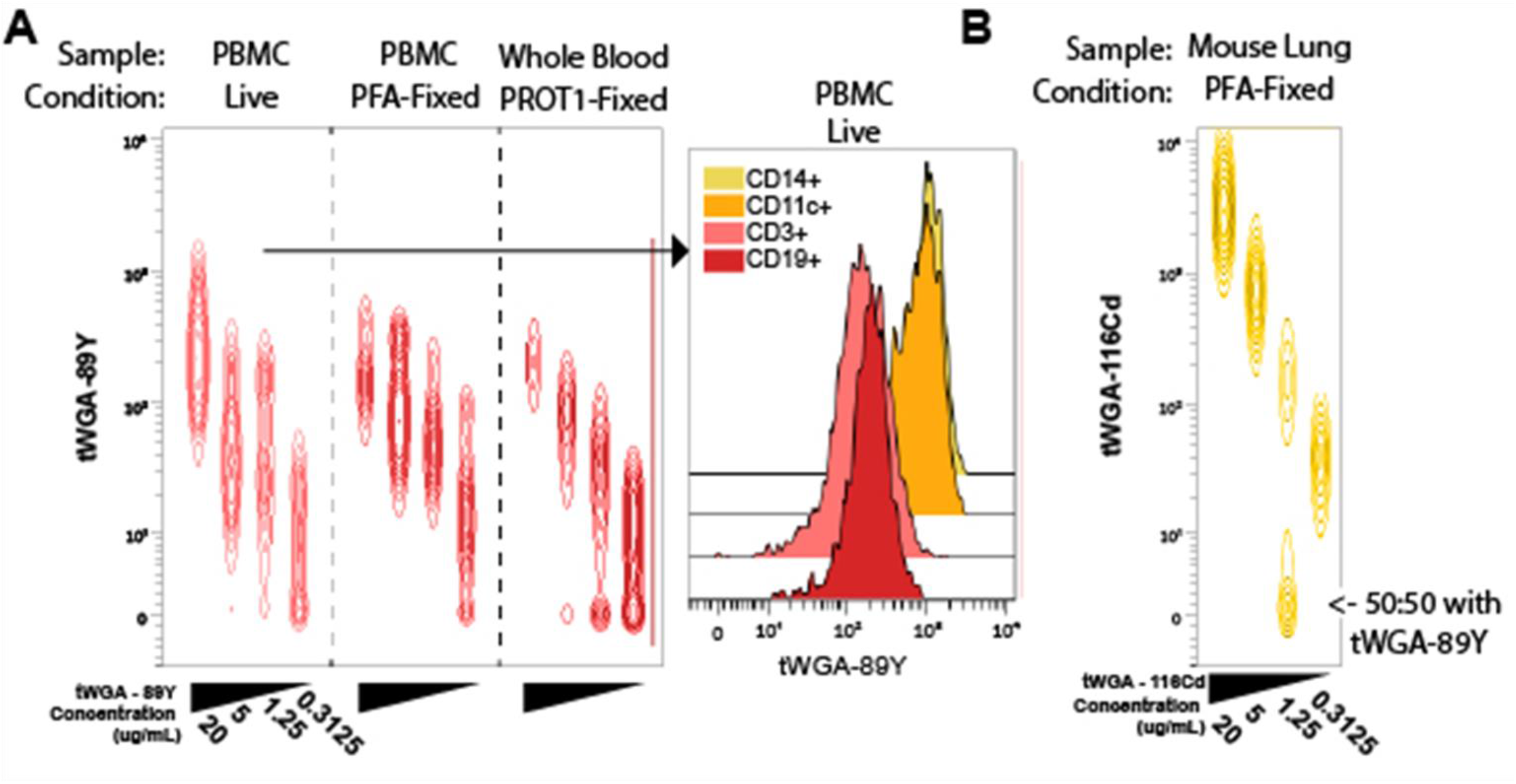
tWGA-metal conjugates stain live and fixed human PBMCs, and fixed mouse lung digest. (A) tWGA conjugated to 89Y was used at a series of titers (20, 5, 1.25, 0.3125 µg/mL) against live, 1.6% paraformaldehyde fixed (PFA-Fixed) and proteomic stabilizer PROT1 fixed human PBMCs. (A, right arrow) Histogram of tWGA-89Y intensity by surface lineage marker from the 20 µg/mL tWGA-89Y titer group. (B) tWGA conjugated to 116Cd was used at a series of titers (20, 5, 1.25, 0.3125 µg/mL) against 1.6% paraformaldehyde fixed (PFA-Fixed) mouse lung digest. The 1.25 µg/mL titer group had an addition of tWGA-89Y stained cells in a 1:1 ratio from the same lung digest. Experiments were run once.

### 20-plex tWGA barcoding on live human PBMCs and fixed mouse lung digest indicates broad applicability of tWGA barcode

In our effort to compare the performance of tWGA barcodes across various samples, we modified our debarcoding workflow introduced previously, by isolating live singlets prior to debarcoding (Figure 4A). Starting with live singlets avoids skewing of performance metrics due to debris, dead cells, or other acellular events which can differ massively between different sample types. Our primary metric for performance is barcode yield, the ratio of all events (in this case, cells) in the source file, to the total number of debarcoded cells which pass barcode QC gating. This hierarchy of cells and simple formula can be seen in Figure 4B. Overlaid debarcoded files from both the PBMC (Figure 4C) and lung digest (Figure 4D) show the typical ‘good debarcode’ lobe above bc_separation_distance of 0.1 and mahalanobis_distance below 20. The mean barcode yields were 0.80 and 0.73 for PBMC and lung digest, respectively. Within live PBMCs, we found no cell-lineage based differences in debarcoding yield (Supplemental Figure 3) Finally, we compare the live-singlet based barcode yield of these two samples against prior assays with available data. These samples using either anti-CD45 live cell barcode or the SBT 20-plex Pd barcode (Figure 4E) on cells of various sources (PBMCs, mouse tissue digests, and cell culture).

**Figure 4:**
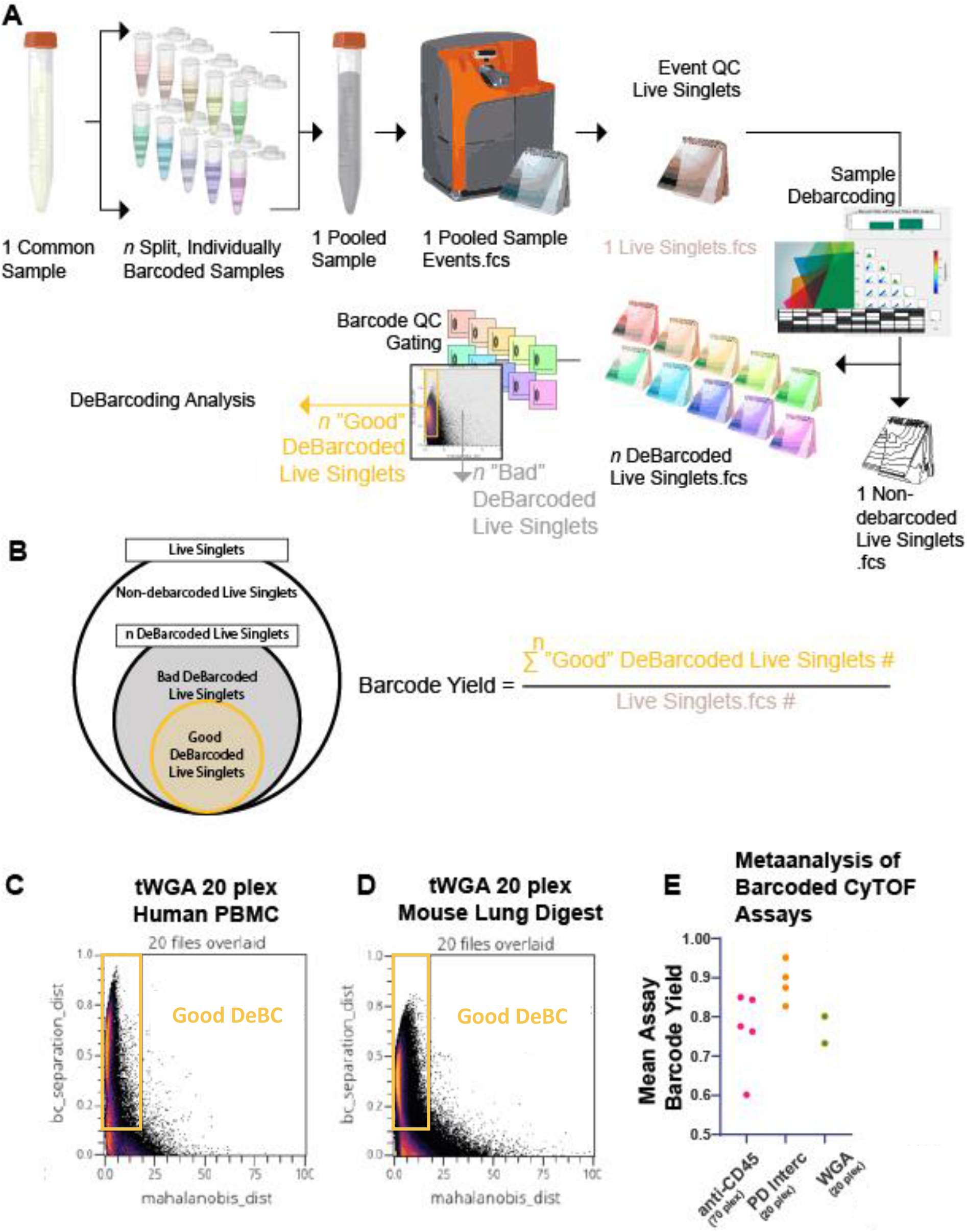
20-plex tWGA barcode analysis workflow and yields comparison. (A) Schematic of experimental design. A single starting sample is split into aliquots for barcoding, each of which receives one of twenty unique barcodes. Post barcoding, samples are pooled and acquired. Output FCS files were gated on live singlets and re-exported. These live singlets are debarcoded and “Good” DeBarcoded live singlets gated on a plot of barcode_separation_distance by mahalanobis_distance. (C) Overlaid density plot of debarcoding QC for the 20 debarcoded PBMC files. (D) Overlaid density plot of debarcoding QC for the 20 debarcoded mouse lung digest files. (E) The per-experiment mean barcode yield from several assays using either anti-human CD45 barcode in an 8-pick-4 arrangement (70 plex), Standard BioTools palladium 6-pick-3 barcode (20 plex), and the aforementioned WGA 6-pick-3 (20 plex) barcode experiments.

### tWGA barcode ‘wandering’ during barcoded sample pooling quantified as barcode yield

Due to the non-covalent, low affinity nature of WGA-ligand binding (micromolar against GlcNAc^26^), we tested whether WGA barcodes were stable in a pooled, barcoded sample prior to fixation. Sample prep is described in Figure 5A. A single sample of live PBMCs was split into ten tubes, five pairs of two. Each pair of tubes were given a pair of complementary barcodes with final effective titers of 20, 10, 5, 2.5, 1.25 µg/mL. These concentrations reflect the total amount of WGA contained within each barcode and not the individual contributions of each conjugate. The complementary barcodes were barcode A (106Cd+110Cd+111Cd), or barcode B (112Cd+113Cd+114Cd+116Cd). This, in theory, generates a ‘worst case’ for barcode mixing in the pooled sample, as any transfer of mass between two barcodes involves mutually exclusive masses. Once the samples were pooled, aliquots were taken at the indicated intervals of 0, 10, 30, and 60 minutes. Samples were then surface stained for 15 minutes and subsequently processed as usual for acquisition. Barcode yields were calculated as described above, and plotted as yield vs pooling time, colored by barcode titer, in Figure 5B. Increases in barcode yield were seen up to ten microgram per milliliter, with twenty micrograms per milliliter exhibiting a decreasing yield with longer pooling times.

**Figure 5:**
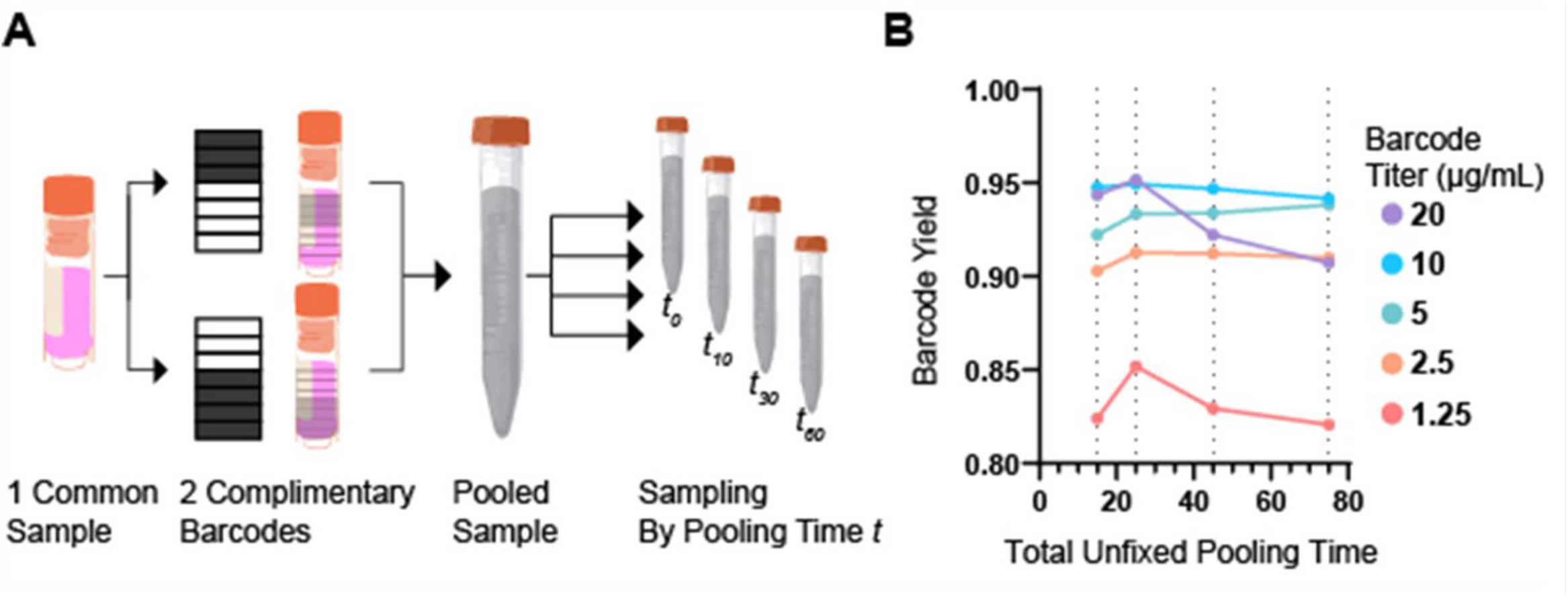
tWGA barcode yield is stable for at least one hour in pooled, barcoded PBMC. (A) Schematic of experimental design. Single sample is split into aliquots. Pairs of aliquots are stained for a titer of a complimentary barcode. Samples are washed and pooled. At the specified pooling time *t*, a fraction of the pooled sample is taken for a surface staining protocol which adds fifteen minutes to the total pooling time. After staining, sample processing proceeded as standard Samples were acquired and subsequently debarcoded. (B) Plot showing barcode yield against pooling time for various titers of barcode. Experiments were run once.

## Discussion

We have reported the first use of Traut’s thiolated WGA (tWGA) as a “plug-and-play” reagent suitable for commercially available maleimide-thiol conjugation kits. 2-iminothiolane (Traut’s Reagent) has been previously used for WGA conjugation in drug delivery applications^27–29^. Traut’s Reagent was chosen to thiolate WGA because the standard reducing preparation for thiol-maleimide conjugations (Tris(2-carboxyethyl)phosphine hydrochloride, or TCEP) resulted in the loss of properly massed WGA as measured by spectrophotometry. tWGA should be compatible with all maleimide-functionalized metal chelating polymers, enabling a broad range of conjugable masses. tWGA retains its membrane-binding properties and can be seen to bind adherent culture cells, live and fixed human and murine cells after conjugation to fluorochrome or metal chelating polymer.

A 20-plex tWGA barcode was created using commercial conjugation kits and used with success on live human PBMC and fixed mouse lung digest. Upon examination of PBMC lineages, we found no cell-type specific differences in barcode yield, which is examined in Supplemental Figure 3. We cannot, however, guarantee tWGA’s functionality under every potential experimental perturbation. An examination of barcode wandering reveals a wide range of “safe” barcode titers, in which the final barcode yield does not decrease due to tWGA or mass dissociation for over an hour. At the highest titer tested (twenty micrograms per milliliter) a progressively decreasing yield is observed with pooling time. This decrease in performance is presumed to be the result of saturation of available cell-surface, increasing tWGA-cell surface turnover in the pooled sample.

The analysis of barcode yields across different reagents, based on re-exported live-singlets, is also novel. Comparing the performance of the myriad combinations of barcode and sample source is not frequent in the literature, and it behooves the community to have a more complete picture of the advantages and disadvantages of these methods. [**note to preprint readers – if you are willing to share non-debarcoded data and a debarcoding key, please reach out! We are hoping to expand the data in the barcode comparison plot in Figure 4! Many thanks to those who already provided data**.].

The choice of isotope conjugates evaluated in this manuscript was driven by a desire for interoperability with extant CyTOF panels using a palladium or cadmium-based barcode. However, we encountered several difficulties with poor conjugate performance which deserve mention. A 20-plex barcode was generated with 115In for the experiments on live PBMC and fixed lung tissue in Figure 4. The dynamic range of tWGA-115 was found to be poor, and all 115In containing barcodes of the PBMC *six-pick-three* 20-plex experiment exhibit poor barcode separation. This poor performance is examined in more detail in Supplemental Figure 4. In our analysis of barcode wandering, our original intent was for an *eight-pick-four* 70-plex barcode. This barcode included tWGA-89Y in addition to the 7 cadmium mass series (106, 110, 111, 112, 113, 114, 116). However, testing of pilot barcodes (1 and 70) indicated the dynamic range of tWGA-89Y was not equivalent to the cadmium series, which lowers the overall performance of any tWGA-89Y containing barcode as a unit. We elected to remove 89Y in our analysis of complementary barcode wandering (Figure 5) as it impacted our ability to resolve the effect of concentration and pooling time on barcoding yields. This “uncounted” tWGA-89Y likely reduces the barcode yield in our experiment vs a true *seven-pick-three* 35-plex barcode, as the tWGA-89Y binding cell surface would preclude the binding of tWGAs conjugated to barcode-relevant masses. Across these assays, the inclusion of poorly performing conjugates 115ln and 89Y into barcodes almost certainly reduces measured barcode yield, and proper adjudication of masses should be expected to increase barcode yields closer to that of SBT Pd barcode levels.

This report represents a proof of concept and merits further optimization. We did not explore using other lectins or combinations of lectins (e.g. concanavalin A), which would presumably increase the available cell-surface ligand to barcode. It is possible that higher levels of Traut’s reagent or longer reaction times during WGA thiolation could further increase the relative labeling performance of tWGA conjugates by increasing available sulfhydryl residues for reaction with maleimide. The recent commercial availability of reactive barcode reagent (Tellurium-Maleimide) may obviate the need for tWGA in live cell barcoding when reaction of cell-surface thiols is acceptable.

Mass cytometry has inherent advantages over fluorescence cytometry when profiling heterogeneous sample. With fluorescence, intrinsic autofluorescence varies between and within cell types, necessitating laborious cell-type- and treatment-specific controls for simultaneous interrogation of multiple cell types^30,31^, while CyTOF generally does not require consideration of cell type for compensation or other corrections. The ability of tWGA barcodes to extend multiplexing to all cells in heterogeneous sample further enhances this advantage. tWGA barcode is unique amongst other affinity reagent in that it seems truly agnostic, barcoding cells regardless of species, lineage, or fixation state. The barcoding and debarcoding process is identical to existing mass cytometry barcoding workflows and requires no protocol modification.

All barcode reagents come with tradeoffs, and no barcoding process will be completely lossless. This is a caveat of all barcodes. Some require chemistries which have a low degree of labeling (DoL) or are too complex or bespoke to be recreated in an average biomedical lab. Affinity reagent requires consideration of inconsistent antigen presentation across cell types or experimental treatments. Reactive reagent comes with concerns of cytotoxicity and the potential chemical modification of antigen, and typically requires more stringent storage conditions than affinity reagent. In short, there is likely no perfect reagent which maximally satisfies all possible criteria. We hope this report of tWGA barcoding provides a useful tool in the sample multiplexing toolbox, for those applications in which it is suited.

## Acknowledgements

Eli Zunder, Corey Williams, Chantel McSkimming for feedback on the project at various stages.

## Materials and Methods

### Generation of tWGA

One milligram of WGA was weighed into a low-bind Eppendorf microfuge tube and reconstituted with 1mL of PBS + 2mM EDTA pH 8.0. Traut’s reagent was reconstituted at two milligrams per milliliter in deionized water. Molar excesses of Traut’s reagent were added as described (0, 2, 5, and 10-fold molar excess of Traut’s reagent to WGA), and the reaction allowed to proceed with mild shaking for at least one hour at room temperature. 7KDa MWCO Zebra spin column was used to exchange the reaction mix for PBS, and the final concentration of thiolated WGA verified by spectrophotometer. Thiolation was confirmed via Ellman’s assay. WGA cartoon derived from published structure ^32^ accessed via the RCSB Protein Data Bank ^33^ and rendered using Mol* viewer^34^. Chemical structures generated in ChemDraw.

### Conjugation of tWGA to AlexaFluor 647 and Fluorescence Imaging

The various thiolations of tWGA were conjugated with AlexaFluor 647 C2 maleimide (Invitrogen A20347) in a 10-fold molar excess for 1.5 hours at room temp. Reaction was washed in a 7KDa MWCO Zebra column to remove unreacted fluor. Human dermal fibroblasts were seeded in a 24 well plate at 60000 cells per well. Cells were stained with each thiolation of tWGA-AF647 mal and with commercial WGA-AF555 (Invitrogen W32464) for 10 minutes at 37c. Cells were acquired on a confocal microscope using appropriate excitation and emission settings for both fluorochromes and mean per-cell integrated fluorescence intensity quantified in Fiji.

### Imaging Flow Cytometry

tWGA 10x Traut’s -AF647-mal was used to stain live human peripheral blood mononuclear cells (PBMCs) at indicated temperatures for 30 minutes. These cells were fixed and counterstained with DAPI prior to acquisition on an Imagestream. Nucleated cells were exported and analyzed. (Cytek, Imagestream X MKII, Inspire and IDEAS 6.2).

### Conjugation of tWGA to metal-chelated polymer

All tWGA conjugations followed manufacturer protocols for antibody conjugation, with the modification that all antibody-reducing steps were omitted and a mass equivalent of tWGA introduced into the prepared, metal-loaded polymer immediately subsequent to the skipped antibody reduction step. 89Y (gift from Dr. Eli Zunder) was conjugated to Maxpar X8 Polymer following manufacturer’s protocol, substituting the provided metal with an equivalent molar amount of 89Y. All other conjugates were derived from Standard BioTools conjugation kits except 115In (IonPath 600115).

### Creation of Barcode

Briefly, stocks of barcode were made from tWGA conjugates as described in the literature^4,7,16^ and diluted to final concentrations as indicated per assay. If not stated, the barcode concentration is ten micrograms per milliliter in a staining volume of fifty microliters.

### Cell Barcoding + Surface Staining

Unless otherwise specified, cell staining protocol is as follows: Cryopreserved PBMCs (90% FBS+10%DMSO) stored in vapor phase LN2 are taken immediately into a 37C water bath, thawed until slushy, and diluted in warm RPMI+10% FBS +10 units/mL bovine pancreas DNAse I (Sigma-Aldritch). Mouse lung digests are derived by 1mg/mL collagenase A (Roche) and DNAse I (50units/mL) digestion + mechanical dissociation of all lobes for 45 minutes at 37c. Digest is then filtered at 100 microns. All subsequent steps proceed at 4c. Cells are spun down at 300xg for 10 minutes, counted and resuspended up to 5E6 cells/mL, and provisioned for sample groups. Cells are then spun down at 300xg for 10 minutes, resuspended in 50uL of cell staining buffer (Standard BioTools or PBS+0.1% BSA) with a unique barcode, and incubated for 30m at 4c. 1mL of staining buffer is added, cells are spun and washed 2x with staining buffer. After washing, cells are then combined into a single 15mL conical which was preadsorbed with staining buffer. Sample is spun down and resuspended in blocking buffer (cell staining buffer + 2.5% FC Block (BD, Fc1.3216) for 15 minutes. Surface-staining antibody panels (including FC block) are made up in a single master mix at 2-5x concentration and frozen at -80c in test-sized aliquots until thawed and added to samples for a final test volume of 100uL. Samples are stained for 30 minutes. Sample is spun, washed and resuspended in a solution of 1:1000 5mM cisplatin (Standard BioTools 201064) for 5 minutes at room temp (final cisplatin concentration 5 uM). Samples are spun and washed twice, and then 16% fresh, EM-grade PFA (Electron Microscopy Sciences) are added to the cell suspensions for a final concentration of 1.6% and allowed to fix for 10 minutes at room temperature. Fixed cells are spun at 800xg for 5 minutes, washed once, and then spun and resuspended in Maxpar fix and perm buffer (Standard BioTools 201067) with 1:1000 of intercalator (Standard BioTools 201192) for 1hr or overnight. Cells are spun and washed 3x in cell acquisition solution (Standard BioTools) prior to acquisition on a Standard BioTools Helios Mass Cytometer.

### CyTOF Normalization, Bead Removal, and Debarcoding

FCS files have EQ beads removed and are normalized using the Nolan lab normalizer^35^. Any experiments acquired across multiple instrument acquisitions are normalized together. Where relevant, samples are debarcoded using the Zunder lab debarcoder^4,5^

### CyTOF Cleanup and Debarcoding QC

Samples are gated using various parameters vs time, with clogs and other acquisition-interrupting events excluded by boolean gating. Event length, width, residual, offset, and center gates are used to further reduce incidence of beads, debris, and doublets. Further bead removal is performed if necessary. Absolute thresholds for barcode separation (no less than 0.1) and mahalanobis distance (no more than 20) are used to ensure no poorly-debarcoded events are carried into subsequent analyses. Beyond those minimums, barcode quality gating is performed manually as described by Fread *et al*^5^.

### Calculation of ‘Barcode Yield’

To compare heterogeneous barcoding datasets, an attempt was made to reach relative parity by isolating and re-exporting live singlets prior to debarcoding. Normalized data are cleaned up as described in ‘CyTOF Cleanup’ above, live singlets gated, and FCS files exported on the live singlet gate. These exported live singlets are then run through the Zunder lab debarcoder with an appropriate sample key. The debarcoded files (including the “unassigned” events file) are then used to generate population statistics for barcode separation (bc_sep) and mahalanobis distance (maha_d). Debarcoding yield is calculated as the ratio of good debarcoded events (sum from all barcodes) vs total events (includes events not debarcoded and events not passing debarcoding QC).

**Supplemental Figure 1.**
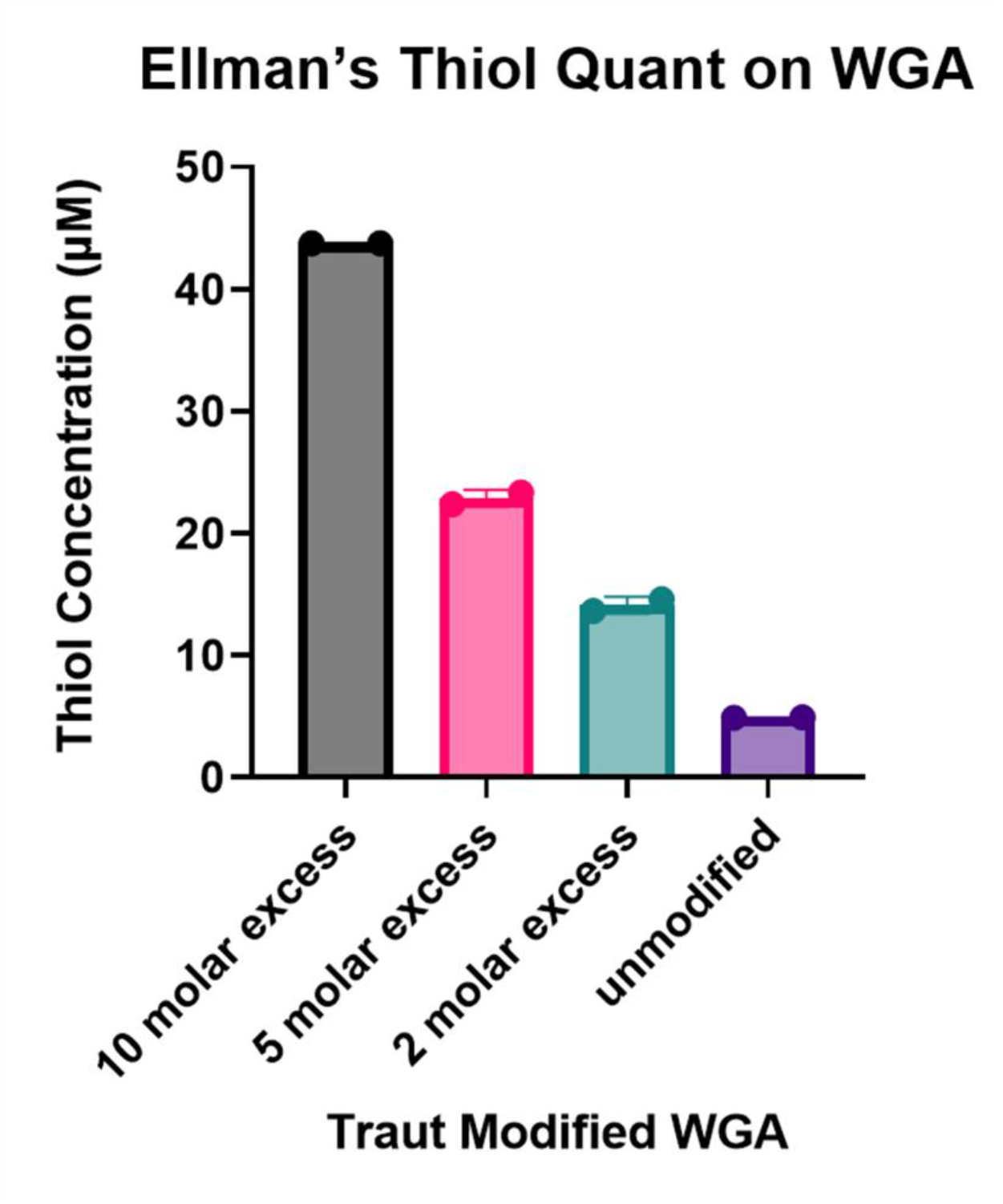
Quantitation of thiolation of Traut’s modified WGA (tWGA) via Ellman’s assay.

**Supplemental Figure 2.**
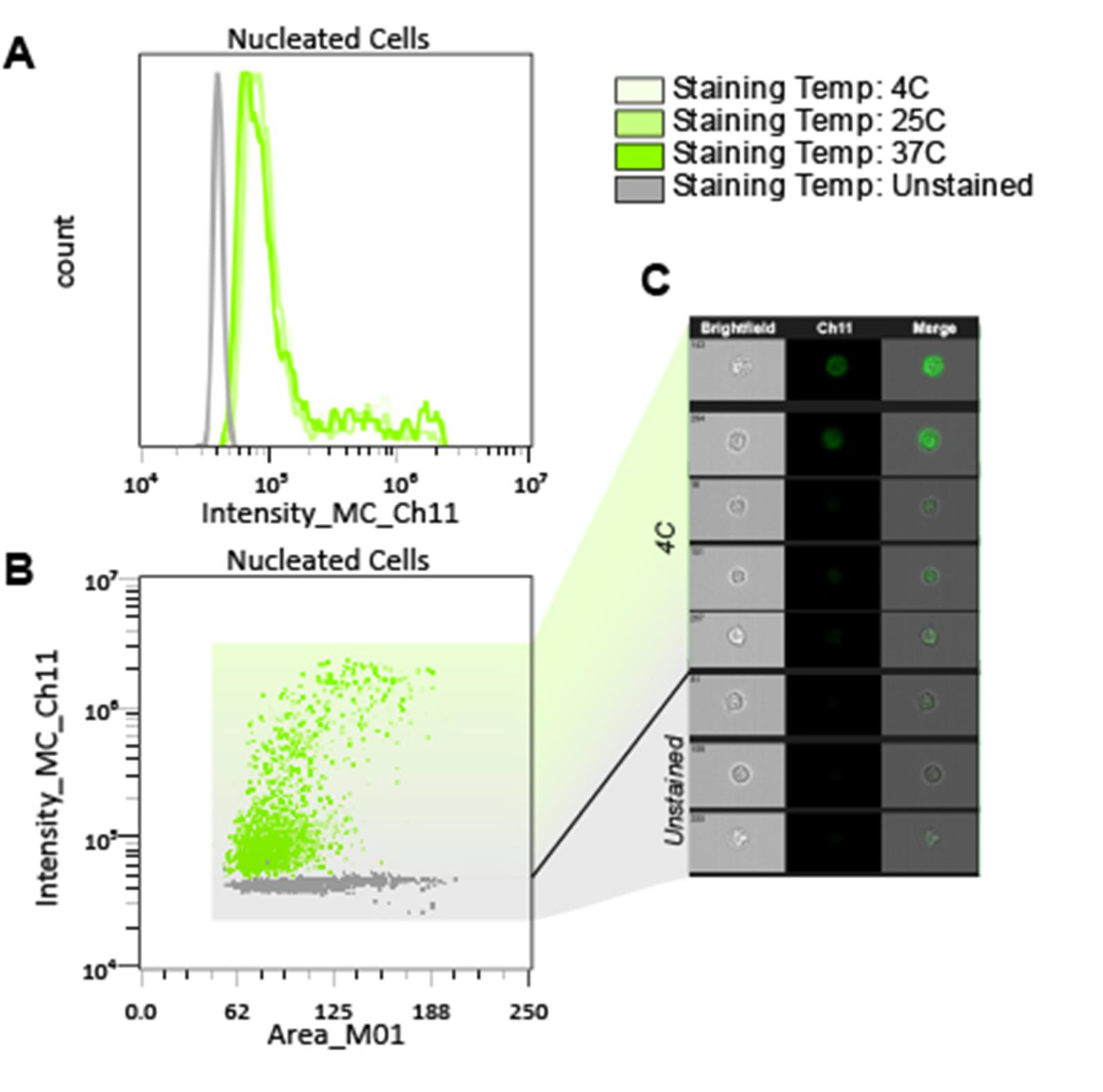
(A) Fluorescence intensities of tWGA-647 stained live PBMCs for 30m at three different temperatures prior to acquisition on an ImageStream imaging cytometer. (B) Biaxial of intensity (y axis) vs area (x axis) of PBMCs. Stained sample’s events colored green, unstained control sample’s events in grey. No significant differences in MFI were seen between the three separate staining temperatures (One way ANOVA, Tukey post-test) (C) Representative images of 4C stained and unstained cells.

**Supplemental Figure 3.**
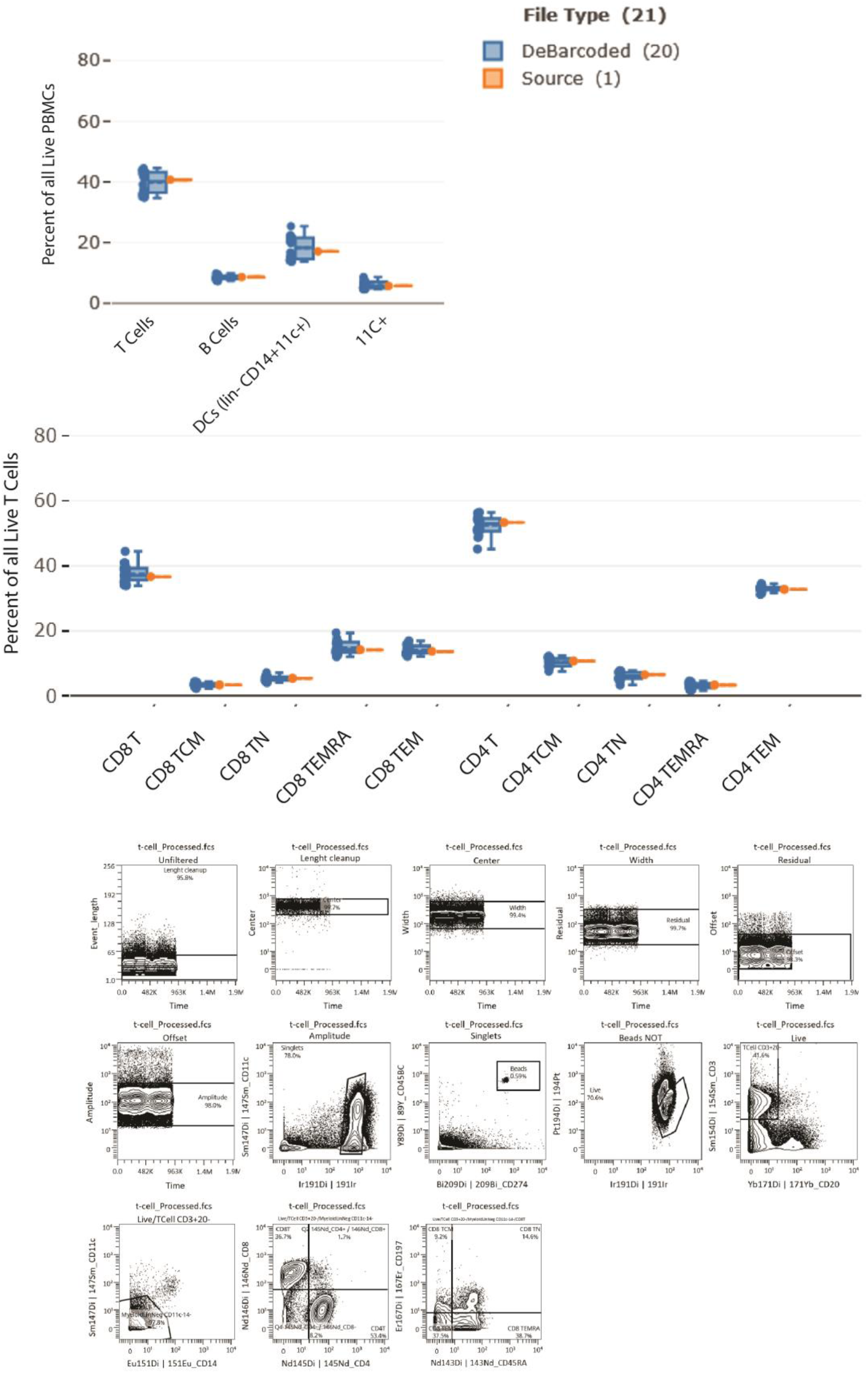
Representative data of human PBMC lineage frequency from either a source file (orange) or the debarcoded files from the source (blue).

**Supplemental Figure 4.**
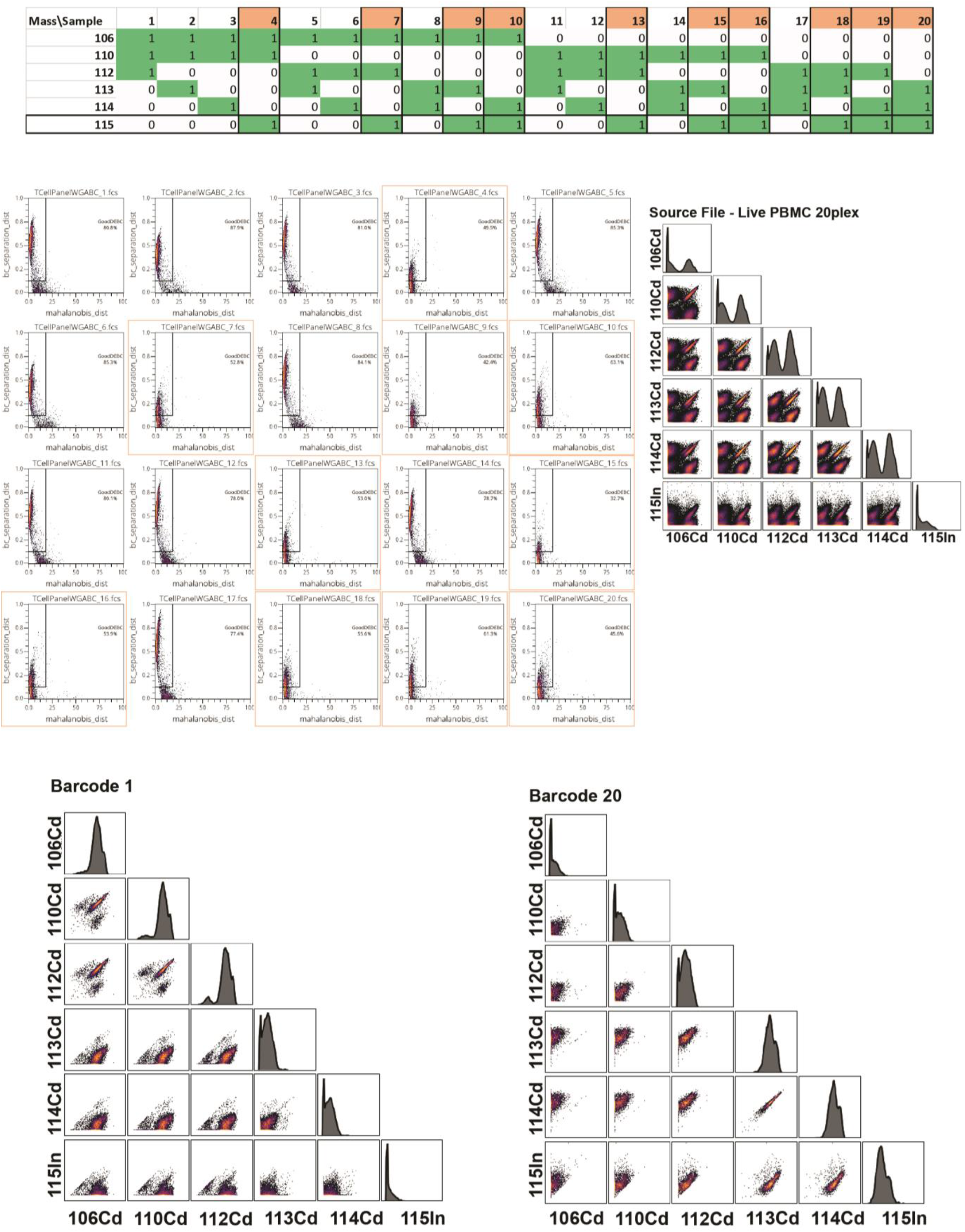
tWGA-In115 barcodes perform poorly in a *six-pick-*three 20-plex barcode of Cadmiums (106, 110, 112, 113) and 115In. Barcodes containing 115In are highlighted in orange.

